# Evolutionary stability of social interaction rules in collective decision-making

**DOI:** 10.1101/2022.12.21.521065

**Authors:** Anna Sigalou, Richard P. Mann

## Abstract

Social animals can use the choices made by other members of their groups as cues in decision making. Individuals must balance the private information they receive from their own sensory cues with the social information provided by observing what others have chosen. These two cues can be integrated using decision making rules, which specify the probability to select one or other options based on the quality and quantity of social and non-social information. Previous empirical work has investigated which decision making rules can replicate the observable features of collective decision making, while other theoretical research has derived forms for decision making rules based on normative assumptions about how rational agents should respond to the available information. Here we explore the performance of one commonly used decision making rule in terms of the expected decision accuracy of individuals employing it. We show that parameters of this model which have typically been treated as independent variables in empirical model-fitting studies obey necessary relationships under the assumption that animals are evolutionarily optimised to their environment. We further investigate whether this decision making model is appropriate to all animal groups by testing its evolutionary stability to invasion by alternative strategies that use social information differently, and show that the likely evolutionary equilibrium of these strategies depends sensitively on the precise nature of group identity among the wider population of animals it is embedded within.

## Introduction

Animal groups exhibit complex collective behaviours that emerge from interactions between social animals [1, 2], such as patterns of collective movement or decision-making. A substantial body of research has established that relatively simple interactions between individuals can produce cohesive groups able to perform complex tasks like self-organised motion [3, 4], group migration [5], conflict resolution [6], or consensus decisions [7]. Substantial effort has been made to identify what these ‘rules of interaction’ are from two perspectives. In one, models are proposed and demonstrated to exhibit the required collective behaviour [3, 8, 9, 4, 6, 10, 11, 12, 13, 14, 15, 16]. This fulfils a necessary but not sufficient condition for identifying the appropriate rules, since other models may also exhibit similar collective behaviour. A second approach is to collect empirical data on animal movements and behaviours and use this to directly infer the form of interactions [17, 18, 19, 20]. Combining these two approaches creates a powerful framework for identifying the rules governing interactions [21].

However, even if one can specify precisely what interactions occur between individuals, this leaves an open question: from among the set of plausible interactions, why do animals use these rules and not others? Although simple interactions between individuals can clearly lead to functional group behaviours, less is known about their evolution and stability on an individual level in groups of unrelated individuals who cannot be assumed to behave according to a single collective goal. Instead, such animals should evolve to make decisions that serve their own selfish interests such as acquiring food and safety. Making informed decisions depends on reliable information about the world. That information comes in the form of cues, which indicate the state of the environment: is there a desired resource over here? Is there a predator over there? As individual sensory abilities are limited, social animals can make use of ‘social information’, i.e. information provided by the actions of their conspecifics, as a source of indirect information about the state of the environment[22, 23]. But not all social information is relevant or accurate [24, 25], and relying too heavily on imitating others can potentially lead to poor information cascades[26]. Therefore we can expect that natural selection will drive animals to adopt specific weightings of private and social information depending on the environment they inhabit so as to maximise the quality of their decisions.

In the area of collective migration, large-scale evolutionary simulations have explored the evolution of interaction rules within a model based on social ‘forces’ [5], with selection on the individual level based on navigational accuracy. This not only demonstrated the evolution of rules sufficient to keep a group of agents together as a single ‘flock’, but also showed the emergence of distinct strategies within the group, characterised by ‘leaders’ and ‘followers’. Importantly, these strategies emerged as a result of individual adaptation under natural selection, rather than being specified in the model itself. Models of collective movement are complex due to the continuous nature of the observable behaviour (motion) and the iterated interactions between individuals over time. As such, it is difficult to make reasoned *a priori* arguments about how animals ought to interact on the move so as to accomplish individual goals, and even evolutionary models such as that above generally work within a heuristic framework of ‘social forces’ – assuming that agents exhibit force-like attraction, repulsion and alignment interactions, and allowing the strength of these forces to be determined by evolution. A more mathematically tractable area of collective behaviour can be considered in the form of simple sequential decision-making between discrete options. Recent research has focused on deriving likely interaction rules in such a scenario by considering the behaviour of rational agents [27, 28, 29]. One such model, developed in refs. [27, 28] has had a considerable influence on empirical work, being used to interpret the observed collective behaviour of fish [30, 31, 32, 33], birds [34, 35] and even humans [36, 37]. However, aspects of this model (described in the next section) remain unspecified by theoretical arguments and must in each case be fitted to the data available. Furthermore, various assumptions made in the model development allow for the possibility that these rules may be vulnerable to exploitation by animals employing a different strategy. Establishing whether the strategy derived in this model is stable is crucial as foundation for the interpretation of the empirical studies which assume its use by the animals under study.

In this paper we take the model of ref. [27] as starting point for considering collective decision making, based on its widespread use in interpreting empirical data. We describe the conceptual and mathematical basis of this model, highlighting potential vulnerabilities due to non-rational assumptions. We identify the key parameters of the model that are left unspecified in theory, and show that these obey necessary relationships under the assumption that animals make decisions optimally, thus reducing the number of degrees of freedom in the model. We then specify alternative strategies an animal might employ using the same conceptual framework, and explore the stability of the baseline model to invasion by these alternatives.

## Method

We consider a commonly used setting of a binary decision [38, 39, 28, 40, 27, 29], where a group of agents needs to decide between two options. One of the options represents the correct decision, be it a safe resting place, the location of a resource etc. The agents choose consecutively, so each focal agent is able to observe two things: the environment and the choices made by the previous agents. The focal agent will then process this available information and use it to make an informed decision about what choice to make.

### Bayesian estimation model

We use a version of the Bayesian estimation model as developed by Perez-Escudero & de Polavieja [27], and we follow their notation and derivation in the description below. The model refers to the above-mentioned setting, and considers each individual making a Bayesian estimation of the probability that each option is the best choice. Each individual estimates the probability that each choice is the best one based on its non-social information *C*, and the behaviour of the other individuals *B*, in order to decide what behaviour to perform.

Let *x,y* be the two available locations the individuals choose from; then, the probability of *y* being the best location is *P*(*Y*|*C*, *B*), and the probability that x is the best location is *P*(*X*|*C*, *B*) = 1 − *P*(*Y*|*C*, *B*). By using Bayes’ theorem,*P*(*Y*|*C*, *B*) is given by:

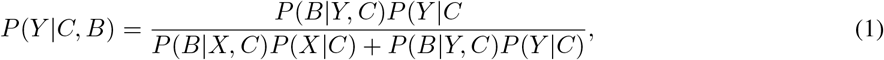

which can be expressed in the following simplified form:

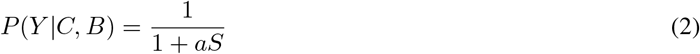

where:

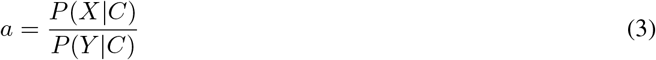

and

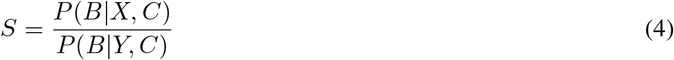

In this model *a* is defined as the non-social parameter as it depends only on the non-social information *C*, while *S* depends on the actions of other individuals. In the supplementary appendices of ref. [27] the authors show that in general *S* may depend on the full ordered sequence of previous decisions. However, in the focal analysis (which has been utilised in the large majority of subsequent studies utilising this model), a simplified version of the model is presented which makes the simplifying assumption that all decisions prior to the focal agent are independent. In this case, *P*(*B*|*X*, *C*)/*P*(*B*|*Y*, *C*) reduces to simple product:

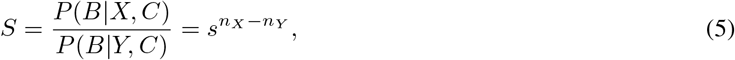

where *s* is a parameter that indicates the relative probability that each agent chooses correctly (i.e. *s* = 2 means that the focal agent assumes each previous decision was twice as likely to be correct as to be wrong). This independence assumption significantly simplifies the form of the calculation, but at the cost of introducing a false belief to the focal agent, which necessarily compromises the optimality of the subsequent decision.

The final aspect of the model relates this Bayesian estimation to the choice the focal agent makes. Following [27], we assume that the focal agent will choose option y with probability *P*(*Y*|*C*, *B*). This assumption allows for the observable reality that an animal confronted with apparently identical conditions and social information may nonetheless make different decisions on different occasions. However, it also introduces a second deviation from rationality, since the probability of choosing correctly is maximised by choosing whichever option has greater than 50% probability of being correct.

### Performance and collective optimality

We now consider how animals employing the above decision strategy will perform in terms of accurately choosing the correct option. To separate this aspect of the model from the conceptual development above, we now consider two options labelled as *A* and *B*, and assume without loss of generality that *A* is correct choice – that is, we take the reward for choosing *A* to be 1 (in some arbitrary units of utility or fitness) and the utility of *B* to be zero. Following the model, each agent chooses either *A* or *B* in turn according to the following probabilistic rule:

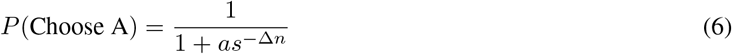

where the non-social parameter *a* defines how reliable the environment is, *s* how strongly the social information is followed, and Δ_*n*_ = *n_A_* − *n_B_* is the social information, specifically the difference in agents that have chosen option A minus the number of agents that have chosen option *B*.

When eq.6 is applied sequentially on all group members on a group with *N* agents, the group will be divided between the two options and will be in one out of *n* + 1 possible configurations, corresponding to the number of agents that have chosen *A*, ranging from 0 to *N*; the probability of each possible configuration depends on the values of *a* and *s*. By summing over the final configurations and their probabilities for a specific set of *a* and *s*, we construct a measure for the group’s collective behaviour, 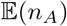, that shows the average number of agents that chose option *A*:

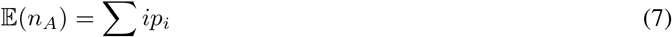

where *i* is the number of agents on option *A* and *p_i_* the probability of i agents being on option *A*. Conceptually we assume that the order of the agents in the sequence is a random permutation for any given decision, such that 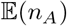 represents the expected reward a randomly chosen agent can expect to receive if all agents apply the same decision-making rule.

Throughout this paper we consider *a* (representing the quality of environmental information) to be a fixed quantity that the agents cannot alter, whereas they may choose a value of *s* to apply. For a given value of *a*, we define the *collectively optimal* value of *s* to be that which maximises the value of 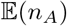.

### Evolutionary stability

So far we have considered the case of identical agents, all of whom make decisions according to a common rule (equation 6), with the same values for parameters *s* and *a*. Under this condition, one can identify a *collectively optimal strategy* that maximises the reward for all agents as above, by maximising equation 7 with respect to *s*; this is the strategy that if employed by all agents of the group, it would lead to the optimal 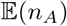 for the group. However, such a strategy is not necessarily evolutionarily stable, since it may be exploited by an individual who applies a different value of *s*. To determine an evolutionary stable strategy (ESS), we must determine a value of *s* = *s*_ESS_ such that if all agents employ this value, no agent can gain by changing their value of to *s* = *s*′.

Here it is important to be precise in how we calculate the effect of an agent varying *s*. In general the expected reward an agent receives for employing a given *s* will depend on its position in the sequence, but we assume throughout that agents do not choose these positions, but are instead randomly shuffled in each decision. Therefore, in calculating the expected reward for an agent employing a new value *s* = *s*′ we average over all the positions in the sequence that this agent might find themselves (with equal probability for each).

Consider a population comprised of identical individuals (all using the same value of *s* = *s*_group_), and one average invading agent using *s*′ = *s_inv_* ≠ *s_group_*. In this case, the average group member will have an expected probability of making a successful choice of 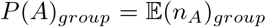, as all agents are identical. The average invader has an expected probability of making a successful choice of *P*(*A*)_*inv*_; this is calculated by using eq.7 for each possible places within the sequence the invader can be in, and taking their average. In this case, the evolutionary stable strategy that the group can employ is the one where no other strategy (i.e. no other value of the parameter *s* = *s*′) can out-perform. The value of *s* where this is achieved is calculated analytically, by considering a range of *s* values, and comparing the rewards for the groups and invader for each one; once these become equal, the respective value of *s* this is occurring for is *s* = *s*_ESS_. As shown in fig.2(a), there is one such value of *s* for the case where both group and invader are using the probabilistic decision rule of equation 6, and it is an equilibrium point.

### Alternative decision rules

Above we consider the evolutionary stability of a given parameter value *s*, assuming that all agents employ the same underlying decision rule specified in equation 6. However, given that the derivation of this decision rule includes multiple departures from full rationality, we anticipate that this could be vulnerable to invasion by alternative decision rules. In particular, adhering to the basic mathematical form of equation 6, two alternatives present themselves as natural variations. In the first case the focal agent does not observe the aggregate number of previous decisions in favour of *A* and *B*, but instead only observes (or responds to) the direction of the majority decision. In this case the appropriate decision rule is:

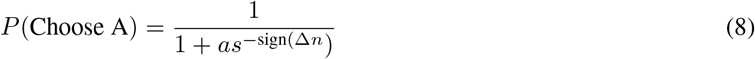

We term this the *simplified* decision rule, since the social information is a lower-dimensional simplification of that in equation 6. This rule is reasonable on two grounds. First, observing the direction of the majority is simpler (especially in large groups) and therefore faster and more reliable as a source of social information. Second, if previous agents had chosen independently (as assumed in the derivation of equation 6), then the Condorcet Jury Theorem implies that the probability that the majority is correct will grow quickly with the number of observed agents.

The second variation we consider is that an agent observes only the most recent decision before its own. Here the decision rule is given as:

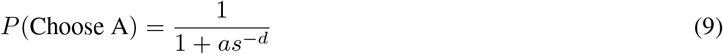

where *d* = 1 if the most recent choice was *A* and *d* = − 1 if *B*. We term this the *dynamic* decision rule in line with similar usage by [32] which investigated an analogous model empirically in humbug damselfish. This rule is motivated by the theoretical findings in the supplementary information of [27] and in [29] that more recent decisions should be weighted more strongly by an agent able to fully account for the correlations in previous agents’ choices, and by the empirical results of [32] and [33] which point to both humbug damselfish and zebrafish responding primarily to the most recent choices of conspecifics.

Similarly to section, we consider a group of agents, where all the agents are using the same decision-making strategy but one average invader who is using one of the other two strategies. Like before, the group and the invader will respectively generate an expected reward 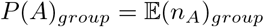 and *P*(*A*)_*inv*_, only now these rewards will depend both on the employed value of *s* and the decision-making strategy, as the strategy determines whether equation 6, 8 or 9 will be used to calculated the probabilistic decision which will in turn generate the values of those probabilities. As before, as long an average invader is able to reach a value of *P*(*A*)_*inv*_ > *P*(*A*)_*group*_ for any value of *s* while using a strategy different to the one used by the group (e.g. the dynamic instead of the aggregate), the group’s strategy is susceptible to invasion. On the other hand, if an invader is not able to outperform the group for any value of *s* while using a strategy other than the one used by the group, then the group’s strategy is stable against the invaders strategy. Moreover, if a strategy employed by the group is not susceptible to invasion by any of the other two strategies for any value of *s*, then that strategy is evolutionary stable; like in section, this stability is calculated analytically.

## Results

### Collectively optimal social behaviour

The performance of a behaviour is measured by the probability of making the correct decision. This depends on the degree of reliability of the environment’s information (value of *a*), and the intensity of following the available social information (value of *s*). Figure 1 shows how variation in the social parameter *s* changes the probability of different group outcomes: panel (a) shows the outcome distribution in the case where *a* = 0.9 and the social parameter is relatively weak (*s* = 1.5). In this case agents are more likely to choose *A* rather than *B*, but intermediate outcomes (those with a roughly equal proportion of agents choosing *A* and *B*) are highly plausible. The probability that all agents will choose *B* is very low. The expected proportion of agents choosing *A* is 0.54092. In panel (b) we show the outcome distribution for the same value of *a* (implying the same quality of non-social information) but a greater value of the social parameter (*s* = 2.3). In this case we make an interesting observation: although the probability that all agents will choose *A* has increased, this has been accompanied by an increase in the probability that all agents will choose *B*, with intermediate outcomes being very unlikely. This has decreased the expected proportion of correct decisions to 0.54029. In panel (c) we show the outcome distribution for the same value of *a* (implying the same quality of non-social information) but a much greater value of the social parameter (*s* = 10). In this case we notice a further increase in the probability that the agents will choose *A* or *B*, and a further decrease in the probability of the intermediate outcomes (with most of them having a 0 probability of occurring). This has decreased the expected proportion of correct decisions to 0.5304. In other words, being more social has increased the probability of making a bad decision. This is due to the probabilistic nature of the system: even with good non-social information, the agents early in the sequence may still make a bad decision. If the tendency to follow social information is very strong, the improbable but still possible bad decision will be copied by the following agents, resulting in an information cascade, eventually misleading a large proportion of the group. This demonstrates that there is a limit to how strongly social information should be followed to maximise collective accuracy.

**Figure 1:**
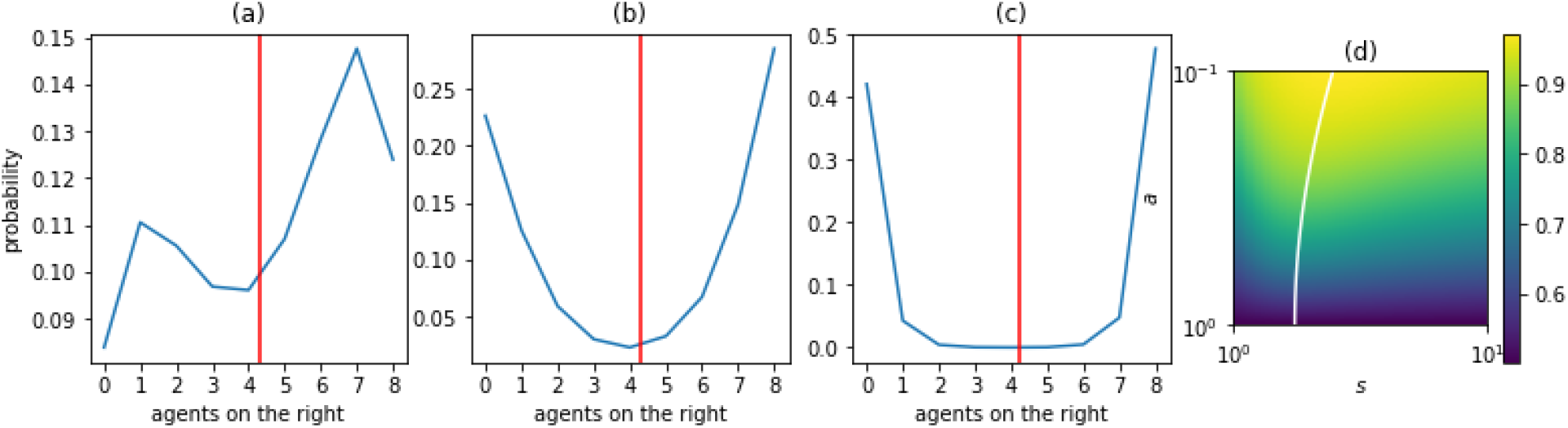
Effect of environmental information (*a*) and intensity of sociality (*s*) on decision-making. Plot (a) shows the probabilities of possible final configurations and the value of 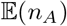 for a group of *n* = 8 agents, *a* < 1 and *s* = 1.5; this corresponds to a case where reliable information is not too strongly followed, leading to very high probability of most agents making a correct decision, and high value of 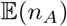. Plot (b) for a group of *n* = 8 agents, *a* < 1 (same as in plot (a)) and *s* = 2.3; this refers to the case where reliable information is followed strongly, now leading to a decrease in most agents making a correct decision and increase to the probability of most agents making a wrong one, and to a slightly lower value of 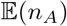 compared to when the same information was followed less strongly. Plot (c) for a group of *n* = 8 agents, *a* < 1 (same as in plot (a)) and *s* = 10; this refers to the case where reliable information is followed very strongly, now leading to a further decrease in most agents making a correct decision (with some cases having a probability 0 of occurring) and further increase to the probability of all agents making a wrong one, and to an even lower value of 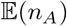 compared to when the same information was followed less strongly. This demonstrates that over-reliance on social information (even when the environment is reliable) can amplify the potentially wrong decisions made by previous agents. Plot (d) summarises the value of *P_R_* for several combinations of *a* and *s*, for *a* ∈[0,1] and *s* ∈[1,10]; we observe higher values of *P_A_* that increase as *a* decreases, while we also observe that across a constant *a*, as *s* increases *P_A_* decreases, as expected due to the aforementioned cost of over-sociality. For *s* > 1 and *a* > 1 (i.e. social behaviour in unreliable environments) we observe a symmetrically opposite behaviour to being social in reliable environments (around the value *a* = 1.)

The effect of *a* and *s* is more widely demonstrated in fig.1(d). This shows the value of 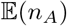 for different combinations these parameters, for *a* ∈[0,1] & *s* ∈[1,10]. We see the observation mentioned previously: being more social in the presence of that information increases the probability of making the best decision up until a point, after which an increase in social behaviour decreases that probability. The white line shown on this panel is the collectively optimal value of *s* for the corresponding value of *a*.

Based on the calculation of 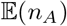 shown in Figure 1(d), it is straightforward to identify the value of *s* that is collectively optimal, shown by the white line. It is clear that as *a* increases (i.e. non-social information becomes less reliable), the collectively optimal value of *s* decreases (agents weight the decisions of others less highly). This makes intuitive sense; as agents are identical, a lower value of a means that other agents are more likely to have made the correct decision, and are therefore more reliable sources of social information.

### Evolutionary stable strategy

Previous research (e.g. [27, 34]) has focused on empirical estimation of *a* and *s* in equation 6 (or the extended version of this model [28]), but estimating both parameters ignores that in a system under natural selection the values of a and *s* should be connected so as to optimise the performance of agents’ decision-making. Here we will determine this necessary connection between the values of *a* and *s* and show that for agents employing equation 6 as a decision-making rule these should not be considered as independent variables.

Above we showed how the collectively optimal value of *s* varies with the reliability of non-social information, *a*. However, this collectively optimal value of *s* indicates the value that would be chosen so as to maximise the success of the group as a whole. As noted in the previous section, under individual natural selection such an optimal value cannot be assumed to be stable (resistant to invasion by other strategies). Instead, we must seek an evolutionarily stable value of *s* = *s*_ESS_ such that a group of agents employing this value cannot be outperformed by an individual who changes their value to an alternative *s* = *s*′. Figure 2 shows the results of this analysis. In panel (a) we show the relative expected rewards for a group employing *s* = *s*_group_ and an invader employing *s* = *s*_invader_ (with non-social parameter *a* = 0.9) – yellow areas show cases where the invader’s reward is greater than the rest of the group, and purple vice versa. As the plot shows, there is a single value of *s*_group_ (indicated by the red line) such that no invader can profit from choosing a different value. This is therefore the evolutionarily stable value of *s*_ESS_ for the particular value of a chosen. Performing this analysis with different non-social parameter values we can map sESS as a function of *a*. This is shown in panel (b) (orange line), alongside the previously calculated value of the collectively optimal *s* (blue line) for comparison. Notably, while both the collectively optimal and ESS values of *s* show a similar pattern of variation with *a* (increasing as non-social information becomes more reliable), they differ markedly across the range of *a* values, with the collectively optimal *s* always being lower than the ESS value. This shows that agents are selfishly motivated to effectively ‘use up’ the available social information, creating strong correlations with other agents that make their own decisions less useful as a source of information to those that follow them. The collective effect of this is to reduce the average performance of all individuals relatively to what they could have achieved had they been able to coordinate on the collectively optimal value of *s*. This ‘price of anarchy’ [reference] (the difference in 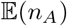 under the two strategies) is shown in panel (c) as a function of *a*, showing a peak at *a* ≃ 0.4.

**Figure 2:**
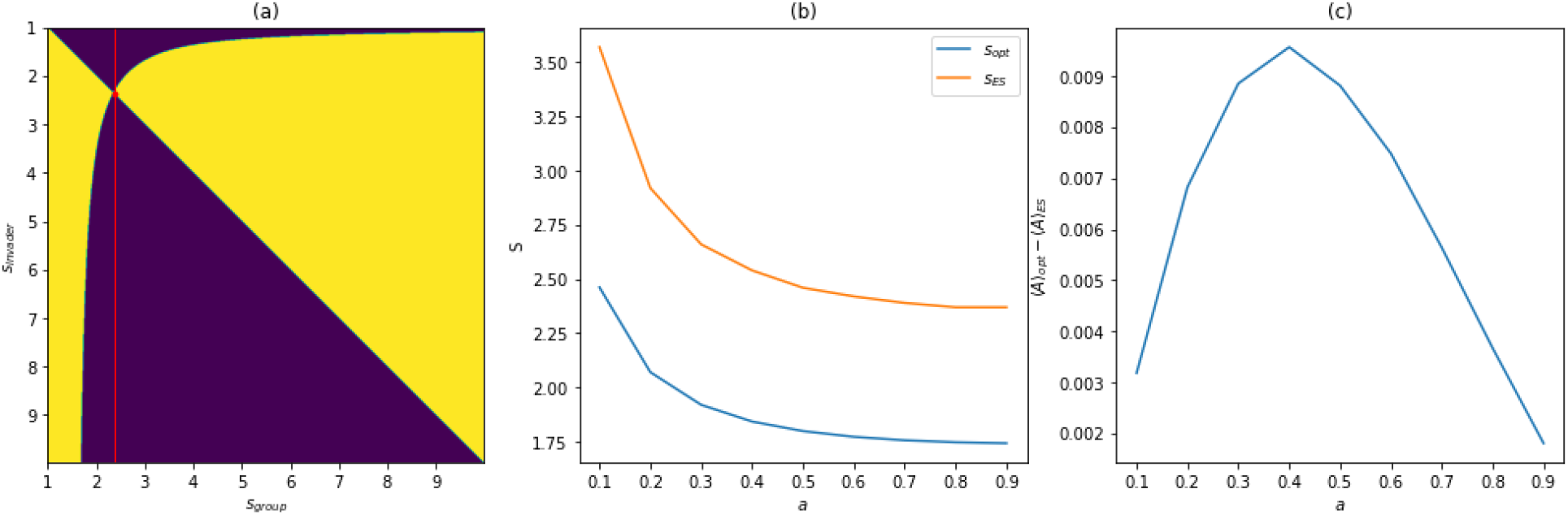
Evolution of social behaviour. Plot (a) shows the dynamics between a homogeneous group and a defector, for different combinations of *s*_group_ and *s*_invader_ and for one value of *a* < 1; the yellow areas correspond to the case where the invader has a higher probability of making the correct decision, the purple areas to the case where the group has a higher probability of making the correct decision, while the diagonal and the curved line correspond to the cases where they have equal probabilities of making the correct decision (with the diagonal being the special case where they actually have the same behaviour, as it is the line where *s*_group_ = *s*_invader_). The intersection of the two lines meets at the evolutionary stable point *s*_ESS_; notice that while the group remains at that value of *s*_group_ = *s*_ESS_, for all values of *S_invader_* the outcome is that the group will have a higher probability of making the correct decision compared to the invader, thus outperforming her. The group reaches eventually reaches that point due to the existence of invaders; in every other point (for all *s*_group_ ≠ *s*_ESS_ the group is outperformed, and will eventually adopt the invader’s *s* as this is more successful-but once *s* = *s*_ESS_ is reached, no other attempt to invade can be successful. Plot (b) shows the values of *s*_ESS_ for the range *a* ∈ [0,1], plotted with the equivalent collectively optimal values *s*_Opt_; *collectively optimal* refers to the value of *s* that the group must use in order to maximise the value of 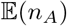 in the environment it navigates, in the absence of invaders. Notice how the evolutionary stable behaviour is not optimal, but over-social. Plot (c) shows how the difference between 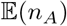 when calculated using *s*_Opt_ and using *s*_ESS_, i.e. the selective pressure the agents in the group our under due to the invader’s presence. We see that the selective pressure is low when uncertainty is low, increases as the uncertainty increases until it reaches a maximum point, and then decreases as uncertainty increases further.

### Alternative decision rules

Animals may observe and respond to social information in a number of different ways. So far we have considered only one decision-making rule, that assumes agents respond to social information in the form Δ_*n*_ = *n_A_* − *n_B_*. We now consider the alternative decision-making rules specified in the Methods section, in terms of the collective behaviour they induce (and whether this is compatible with observations of real animal groups) and their relative performance in decision-making accuracy.

A common characteristic of group decisions is the tendency towards consensus decision making – outcomes in which all agents choose the same option are the most probable [7]. All three decision rules we have tested are able to replicate this collective pattern, as shown in Figure 3, by selection of appropriate values of *a* and *s*. This is prima facie evidence that all three models are suitable candidates for modelling collective decisions.

**Figure 3:**
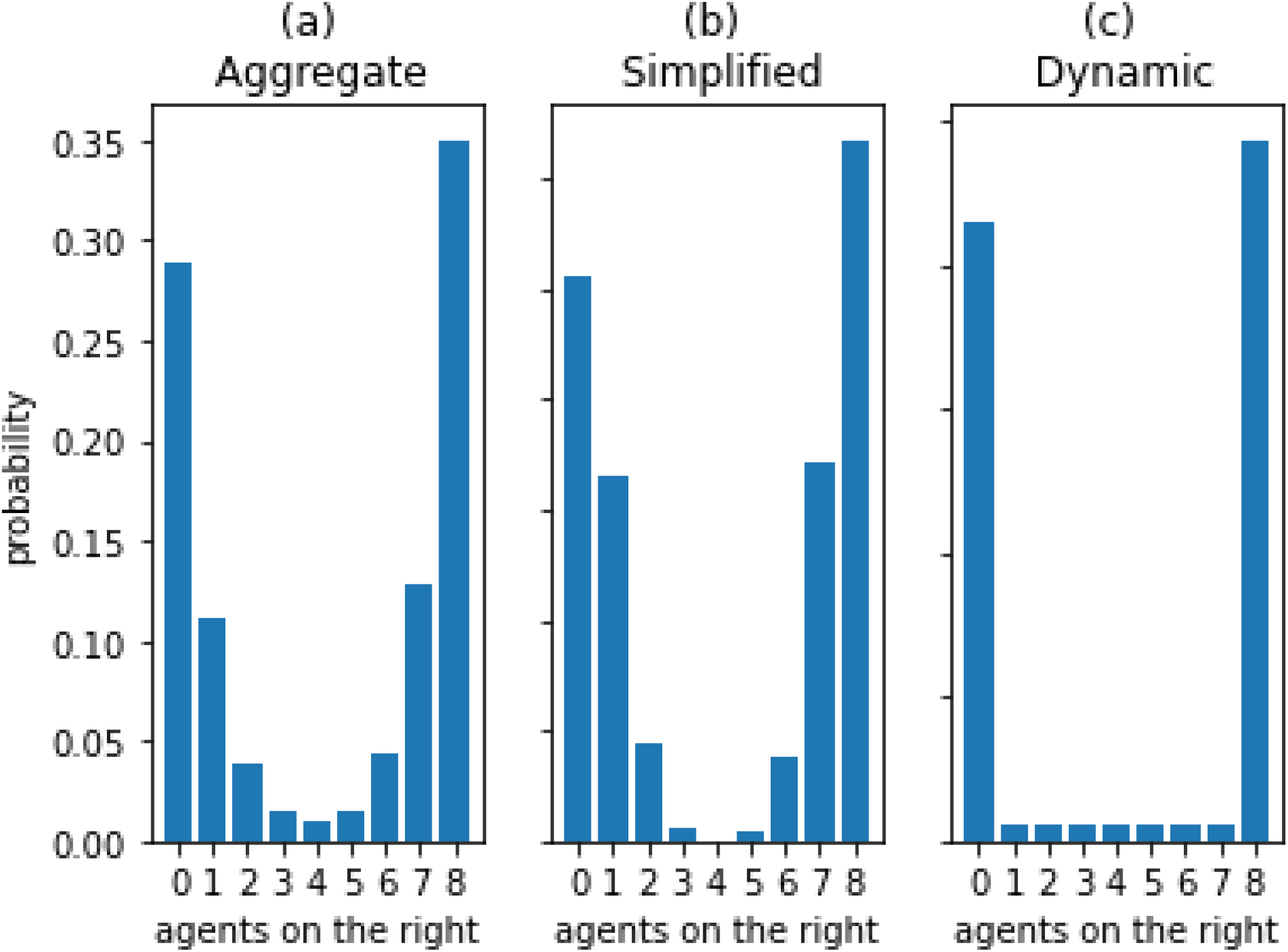
Probabilities of final states, for a group of *n* = 8 agents and different strategies. Each plot has a different pair of *a*, *s*, leading to the same value of 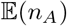 for all strategies and a bias towards consensus-reaching where the probability that all of the agents will make the same choice is high. This demonstrates that we cannot necessarily infer which strategy is being used by the agents simply by noticing that there is a bias towards consensus.

Instead we can appeal to the theoretical performance of each decision making rule as a basis for judging its credibility as a candidate for real animal groups. In our analysis above we determined the evolutionarily stable value of *s* for a group in which agents all employ the decision making rule in equation 6, by analysing whether an invading strategy with a different value of *s* could outperform the other members of the group. We can extend this stability analysis to ask whether an invading strategy with a different value of *s* and *a* different decision making rule can outperform an otherwise homogeneous group.

In fig.4(a) we consider the dynamics between all possible combinations of strategies between group and invader for a group like the one consider before. Along the diagonal are the cases where they both employ the same strategy, while the rest corresponds to the cases where group and invader employ different decision rules. Each column refers to the group using the same strategy (aggregate, simplified and dynamic starting from the left), and every row to the defector using the same strategy (aggregate, simplified and dynamic starting from the top). The yellow areas signal the cases where the invader outperforms the group, while the purple ones the cases where the group outperforms the invader.

**Figure 4:**
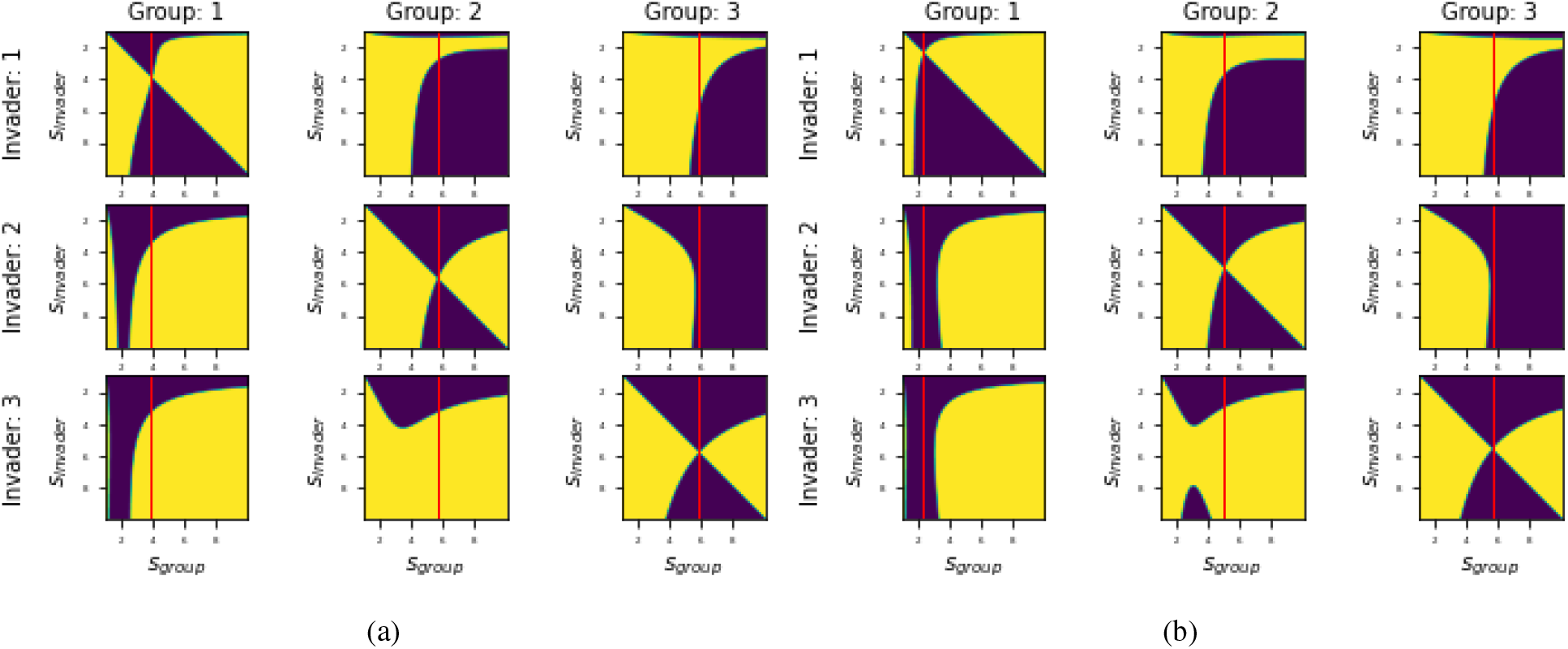
Dynamics between group and invader, for different combinations of decision-making strategy use and *N* = 8, *a* = 0.9. Plot (a) shows the dynamics in the case of a fixed group; here, the invader is assumed to always be present and disrupting the information provided to her peers. In the cases where group and invader employ the same strategy, they are always able to reach an equilibrium point corresponding to the evolutionary stable state of the group; the value of *s*_ESS_ is marked in all three cases with a red line. In each column, this indicates whether a strategy is successful or not against the group’s strategy; this happens whenever that line falls into yellow areas. We notice that for all the strategies, an invader is able to invade using a different one. Specifically: a) the aggregate strategy can be invaded by both the simplified and the dynamic for a wide range or *s* values (ranging from fairly small to large values), b) the simplified strategy can be invaded for a wide range or *s* values by the dynamic strategy (ranging from fairly small to large values) and for a narrow range of small *s* values by the aggregate, and c) the dynamic strategy can be invaded by a range of *s* values (ranging from small to medium). This means that a fixed group does not evolve to use a single strategy. Plot (b) shows the case of an infinite group; in that case, the invader may not always be present during the decision-making process and hence her presence is not as disruptive. In this case we notice that although the simplified and the dynamics strategies can be invaded like before (i.e. the simplified can be invad, the aggregate cannot, meaning that an infinite group can evolve to employ a single strategy (namely the aggregate strategy).

For each decision rule in isolation there exists a single evolutionarily stable value of *s*. Groups employing a single decision rule can therefore be assumed to reach this stable point, where they cannot be outperformed by invaders using different values of *s*. However, they may be outperformed by invaders using a different strategy against it. In each column, the vertical red line signifies the group’s evolutionary stable value of *s*. If this line falls within a yellow region for a different decision rule, it signifies that in that case the invader can employ this different decision rule, with the corresponding values of *s* that are within that region, to outperform the group.

For example, consider the left column of fig.4(a); the top plot shows the equilibrium point for a group employing the aggregate decision rule (equation 6) and the invader’s failure to out-compete using the same decision rule (since the vertical line falls exclusively in purple areas). However, if the defector chooses to employ the simplified strategy (equation 8), as shown in the middle plot, the same line passes through yellow areas, meaning that there are values of *s* the defector can employ to outperform the rest of the group. This implies that the aggregate decision rule is not globally stable against invasion by the simplified decision rule (assuming that invasions can arise freely on any alternative rule and with any value of *s*, rather than being restricted to local mutations)

This does not imply that the simplified decision rule is a stable strategy for the group to employ. Similar inspection of the results as above shows that this rule can be invaded by both the aggregate and dynamic rules. Instead, what this analysis shows is that in a group restricted to employ these three rules, no single decision rule is globally stable. This may eventually lead to the coexistence of different rules in the group, a cyclical transition between rules or the adoption of new rules not tested here. Which of these occurs depends on further assumptions about the evolutionary process beyond the initial invasion, which we do not consider further here. What we can establish from these results is that the aggregate decision rule, which has been widely used as a model for interpreting collective behaviour in real systems, is not stable under the conditions we have described.

So far we have considered a fixed group in which the same agents repeatedly make decisions together. However, many animal groups in which decisions are made are transitory, being drawn from a larger population by (for example) fission-fusion dynamics. In such populations the effect of an invader may be different than in a fixed group because each agent in the population encounters the invader more rarely (the invader is rarely part of any randomly selected subgroup), and thus the majority of an agent’s rewards are obtained in interactions solely with the dominant phenotype. In very large populations the effect of the invader on other agents will be negligible, whereas in a single fixed group the invader may severely disrupt other agents’ use of social information across the whole group.

We investigated whether this changed the stability relationships between different decision rules, assuming that in each decision a group of 8 is drawn randomly from an effectively infinite population. In this case we observe different dynamics between group and invader; as shown in fig.4(b), while in this case the simplified and the dynamic strategy can still be invaded from the other two strategies for some values of *s*, the aggregate strategy can’t be invaded by either. So in that case, eventually, a larger group will evolve to use the aggregate strategy as this is evolutionary stable, under the assumption that only these three decision rules are available.

So far we have considered a group of size *N* = 8 in an environment of *a* = 0.9. Different group sizes lead to different dynamics, since group size affects the information transfer process, and the quality of available social information to the agents. For a fixed value of *a*, different group sizes (*N*) lead to different dynamics; a decrease in group size leads to the group’s strategy being more robust against invaders, as it can be invaded by them for a narrower range of *s* values. Cases with different group sizes and different values of a are briefly presented in the supplementary material.

## Discussion

Collective decision-making emerges from the individuals’ decisions, which in turn are affected by the available personal and social information. Here, we use a probabilistic sequential decision-making model, to understand how different rules of interaction between individuals affect the quality of decision-making. We established strategies for the use of social information that are either optimal for the group as a whole, or evolutionarily stable to invasion by alternative strategies.

The probability of an agent making a good decision depends both on the reliability of the available information, and on how strongly this is followed. While in general a more certain environment means that following social information is beneficial, the probabilistic nature and the non-linear form of the decision-making rule, mean that an increase in social following is not always beneficial. Although social information is valuable, the tendency of agents to follow each other risks magnifying the effect of early incorrect decisions. Thus the collective performance of a group is maximised at some finite value of *s*, indicating that social information is neither ignored nor followed deterministically. As the quality of non-social information increases, so does the collectively optimal value of the social weighting, since the social information provided by other agents increases in value too.

Collectively optimal strategies are vulnerable to exploitation in groups of selfish agents, and here the expected value of social weighting that real groups will exhibit is that represented by an evolutionarily stable strategy (ESS) – one that cannot be successfully exploited by an invading alternative strategy. We find that such an ESS always weights social information more highly than would be collectively optimal. This limits the accuracy of the group, making individual decisions less often correct than they would be if the group could coordinate on the collectively optimal social weighting. This result mirrors similar findings in a related system [41], where the authors conclude that animals will eventually evolve to a ‘sub-optimal’ state of over-reliance on social information. The recurrence of that phenomenon here suggests that such overweighting of social information (relative to what would be collectively optimal) may be a general property of animal groups.

Extending on such previous research as ref. [41], which consider the evolution of social weighting parameters within a single decision rule, we also investigated the evolutionary stability of different decision rules. Here we found our results depend sensitively on our assumptions about the nature of the population the decision-making group is drawn from. When agents are drawn randomly from a large population each time they form a group to make a decision, the aggregate decision strategy derived by ref. [27] is stable to invasion by either of the alternatives we considered. However, if the population of agents is restricted solely to the individuals within a single decision making group, this stability is broken, with all three decision rules potentially coexisting or cyclically replacing each other. These results are limited to the particular decision rules we have considered, and others we have not tested may be favoured by natural selection. Furthermore, real groups likely exist somewhere between the two extremes we have considered, such as in fission-fusion dynamics where small groups within a larger population may remain together for some time. Nonetheless, our results suggest that careful consideration should be given to the nature and stability of group membership when considering whether a particular decision rule (such as equation 6) is an appropriate model for the system under examination.

In our model there is no additional cost to using a more computationally or time consuming strategy. It is likely that counting exactly how many other animals have chosen each option is more costly in these measures than either observing the majority choice, or just the most recent decision. We expect that adding costs to each model will change how group-invader dynamics play out, but determining exactly what these costs are would require specific knowledge of the sensory environment and the factors influencing the cost of time in decision making, so we have have chosen to omit further exploration of these costs here.

We have also chosen to define group behaviour as the averaged behaviour of the group-members. While this is informative as it led to insights about behaviour evolution, this approach ignores the nuance of information within a group; depending on the rank, agents have access to different information, and it’s sensible to expect this to contribute to different behaviours for agents, depending on their place in the group. Furthermore, we expect this to affect the dynamics between group and invader, since an invader’s success depends on what place within the group she manages to place herself. We believe that by including this level of complexity, we will be able to add some insight into the existing literature that researches dynamics between agents with differing (or even conflicting) goals within groups[42, 43, 44].

## Acknowledgments

This work was funded by UK Research and Innovation Future Leaders Fellowship MR/S032525/1 and a PhD scholarship from the University of Leeds.

