## Supplementary Material for "Evolutionary stability of social interaction rules in collective decision-making"

This document provides some supplementary material for the paper 'Evolutionary stability of social interaction rules in collective decision-making', regarding the effect of group size and value of parameter $a$ on the dynamics between a group and an invader employing different strategies.

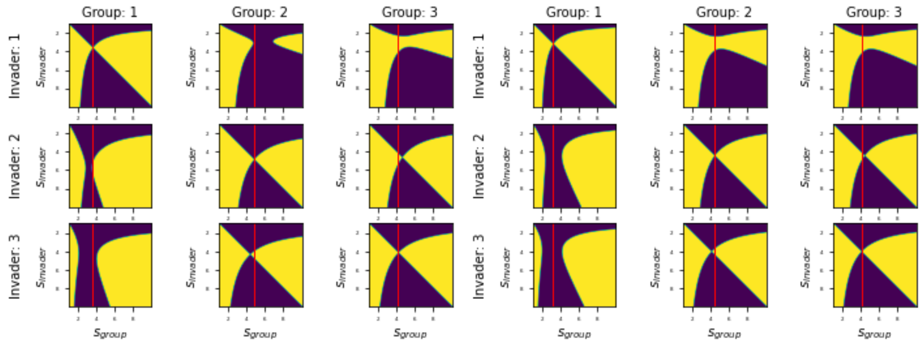


Figure 1: Dynamics between group and invader, for different combinations of decision-making strategy use where $N = 3$, $a = 0.3$. Plot (a) shows the dynamics in the case of a fixed group; the value of $s_{\text{ESS}}$ is marked in all three cases with a red line. In each column, this indicates whether a strategy is successful or not against the group's strategy; this happens whenever that line falls into yellow areas. We notice that for all the strategies, an invader is able to invade using a different one for a narrow range of $s$ values (specifically, the aggregate strategy can be invaded by the dynamic, the simplified can be invaded by the dynamic, and the dynamic can be invaded by both the aggregate and the simplified), meaning that a fixed group does not evolve to use a single strategy. Plot (b) shows the case of a larger group; in this case we notice that although the simplified and the dynamics strategies can be invaded like before (i.e. the simplified can be invaded by both the aggregate and the dynamic, and the dynamic can be invaded by the aggregate), the aggregate cannot, meaning that an infinite group can evolve to employ a single strategy (namely the aggregate strategy).

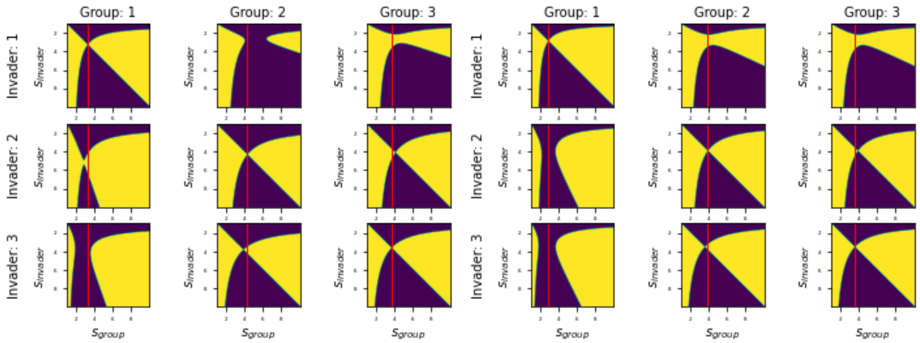


Figure 2: Dynamics between group and invader, for different combinations of decision-making strategy use where $N = 3$, $a = 0.9$. Plot (a) shows the dynamics in the case of a fixed group; the value of $s_{\mathrm{ESS}}$ is marked in all three cases with a red line. In each column, this indicates whether a strategy is successful or not against the group's strategy; this happens whenever that line falls into yellow areas. We notice that for all the strategies, an invader is able to invade using a different one for a narrow range of $s$ values (specifically, the aggregate strategy can be invaded by the simplified, the simplified can be invaded by the dynamic, and the dynamic can be invaded by both the aggregate and the simplified), meaning that a fixed group does not evolve to use a single strategy. Plot (b) shows the case of a larger group; in this case we notice that although the simplified and the dynamics strategies can be invaded, in a different way than before (i.e. the simplified can be invaded by both the aggregate and the dynamic, and the dynamic can be invaded by the aggregate), the aggregate cannot by invaded by any, meaning that an infinite group can evolve to employ a single strategy (namely the aggregate strategy).

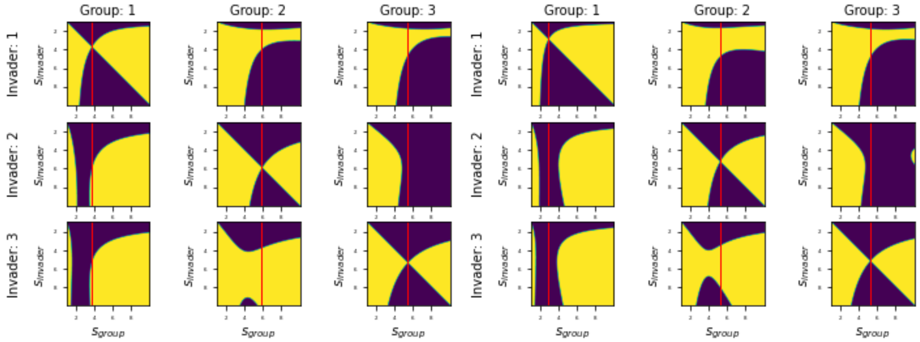


Figure 3: Dynamics between group and invader, for different combinations of decision-making strategy use where $N = 5$, $a = 0.3$. Plot (a) shows the dynamics in the case of a fixed group; the value of $s_{\text{ESS}}$ is marked in all three cases with a red line. In each column, this indicates whether a strategy is successful or not against the group's strategy; this happens whenever that line falls into yellow areas. We notice that for all the strategies, an invader is able to invade using a different one (specifically, the aggregate strategy can be invaded by both the simplified and the dynamic, the simplified can be invaded by both the aggregate and and dynamic, and the dynamic can be invaded by the aggregate), meaning that a fixed group does not evolve to use a single strategy. Plot (b) shows the case of a larger group; in this case we notice that although the simplified and the dynamics strategies can be invaded like before (i.e. the simplified can be invaded by both the aggregate and the dynamic, and the dynamic can be invaded by the aggregate), the aggregate cannot, meaning that an infinite group can evolve to employ a single strategy (namely the aggregate strategy).

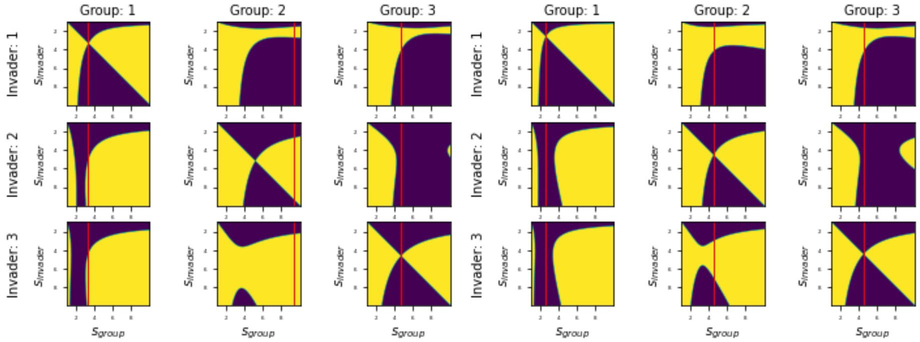


Figure 4: Dynamics between group and invader, for different combinations of decision-making strategy use where $N = 5$, $a = 0.9$. Plot (a) shows the dynamics in the case of a fixed group; the value of $s_{\text{ESS}}$ is marked in all three cases with a red line. In each column, this indicates whether a strategy is successful or not against the group's strategy; this happens whenever that line falls into yellow areas. We notice that for all the strategies, an invader is able to invade using a different one (specifically, the aggregate strategy can be invaded by both the simplified and the dynamic, the simplified can be invaded by both the aggregate and and dynamic, and the dynamic can be invaded by the aggregate), meaning that a fixed group does not evolve to use a single strategy. Plot (b) shows the case of a larger group; in this case we notice that although the simplified and the dynamics strategies can be invaded like before (i.e. the simplified can be invaded by both the aggregate and the dynamic, and the dynamic can be invaded by the aggregate), the aggregate cannot, meaning that an infinite group can evolve to employ a single strategy (namely the aggregate strategy).

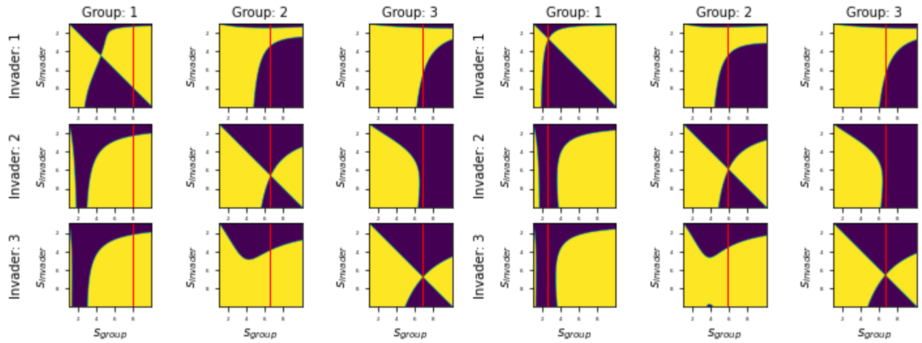


Figure 5: Dynamics between group and invader, for different combinations of decision-making strategy use where $N = 8$, $a = 0.3$. Plot (a) shows the dynamics in the case of a fixed group; the value of $s_{\mathrm{ESS}}$ is marked in all three cases with a red line. In each column, this indicates whether a strategy is successful or not against the group's strategy; this happens whenever that line falls into yellow areas. We notice that for all the strategies, an invader is able to invade using a different one for a wide range of $s$ values (specifically, the aggregate strategy can be invaded by both the simplified and the dynamic, the simplified can be invaded by both the aggregate and and dynamic, and the dynamic can be invaded by the aggregate), meaning that a fixed group does not evolve to use a single strategy. Plot (b) shows the case of a larger group; in this case we notice that although the simplified and the dynamics strategies can be invaded like before (i.e. the simplified can be invaded by both the aggregate and the dynamic, and the dynamic can be invaded by the aggregate), the aggregate cannot, meaning that an infinite group can evolve to employ a single strategy (namely the aggregate strategy).
